# Intranasal Leukemia Inhibitory Factor as a late-stage treatment for delayed white matter damage in concussive head injury

**DOI:** 10.1101/2025.04.07.647435

**Authors:** Veera D’Mello, Jelena Mihailovic, Sidra Ali, Basavaraju G. Sanganahalli, Daniel Coman, Fahmeed Hyder, Merisha Fernando, Anita Mampilly, Sridhar S. Kannurpatti, Steven W. Levison

**Affiliations:** Rutgers-New Jersey Medical School, Newark, NJ, 07103 USA; Yale University, Magnetic Resonance Research Center, New Haven, CT, 06520 USA

**Keywords:** traumatic brain injury, astrogliosis, axonal damage, magnetic resonance imaging, diffusion tensor imaging, corpus callosum, prefrontal cortex

## Abstract

Leukemia Inhibitory Factor (LIF) is an injury-induced cytokine that peaks 48 hours after a traumatic brain injury (TBI). Juvenile LIF haplodeficient mice exhibit desynchronized glial responses, increased neurodegeneration, decreased axonal conductivity and behavioral deficits after a concussive head injury. Given the necessity of LIF during the acute recovery phase after injury, we hypothesized that intranasal LIF (IN-LIF) treatment would prevent neurodegeneration when administered during the chronic recovery period from a mild TBI (mTBI). Young adult male CD1 mice were subjected to a midline, closed-head frontal cortex injury using a flat metal impactor with a 3mm tip to induce a mTBI. In the 6-8 weeks post-mTBI, known to precede axonal atrophy in this mTBI model, two doses of 40 ng and 100 ng of LIF were administered twice daily, 5 days/week for two consecutive weeks. Sensorimotor functions were assessed at 4 and 8 weeks post mTBI, followed by ex-vivo brain magnetic resonance imaging at 9.4T and histopathology. mTBI mice showed sensorimotor deficits at 4 weeks, which worsened by 8 weeks post-injury. IN-LIF treatment prevented the progressive sensorimotor loss seen in the vehicle-treated controls. Increased mean diffusivity and decreased fractional anisotropy were observed in the corpus callosum and prefrontal cortex of mTBI brains. In a dose-dependent manner, IN-LIF prevented the mTBI-induced mean diffusivity increase and fractional anisotropy decrease. Histologically, there was significantly less astrogliosis, microgliosis and axonal injury in the IN-LIF treated mice vs. controls. These results support the therapeutic potential of IN-LIF to reduce delayed neurodegeneration and improve neurological outcomes after mTBIs.

## Introduction

Almost 80% of all traumatic brain injuries (TBIs) are classified as mild (mTBI) based on clinical severity. Whereas it had been thought that a single concussive hit to the head did not produce long-term effects, approximately 1/5 of mTBI patients report poor memory, reduced processing speed, fatigue, impulsivity, anxiety or depression that may be due to delayed neurodegeneration (1, 2). Recently, CTE pathology was found in young (∼23-year-old) deceased athletes, underscoring the fact that unaddressed TBI can progress to more serious neurodegenerative pathologies (3). Analogously, several studies have shown persistent behavioral deficits in single-hit adult mouse mTBI models across various behavioral tasks, including spatial memory (4), olfactory discrimination (5), object recognition (6), passive avoidance (7), the forced swim test (7), and analyses of circadian rhythms (8). Progress is being made in developing neuroprotective strategies to reduce acute brain injury, and several FDA-approved drugs are being investigated (9); however, clinical trials are still in the early stages. By contrast, drugs are lacking that can be administered during the chronic stage of recovery. Indeed, in a clinical trial to evaluate the efficacy of minocycline, microglial activation appeared to be inhibited. However, this did not prevent tertiary neurodegeneration as revealed by increased plasma neurofilament light chain (10) levels.

Histopathological and electrophysiological analyses in a single hit closed head injury of mTBI have provided insights into the chronic phase of neurodegeneration(11). Notably, with this injury, the subcortical white matter (WM) appears normal up to 6 weeks after the mTBI. However, tertiary neurodegeneration becomes apparent after 6 weeks of recovery, where the compound action potential becomes weak, accompanied by Wallerian degeneration (12). Moreover, the enzyme Sterile alpha and TIR motif containing 1 (SARM1), which is the executor for Wallerian axonal degeneration, is necessary for this delayed axonal injury, as the axonal degeneration does not occur in SARM1 null mice after a single hit TBI (13).

Intranasal delivery of drugs into the brain has great clinical potential due to (1) simplicity of administration, (2) noninvasiveness, (3) relatively rapid CNS delivery, (4) ability to repeat dosing easily, (5) non-necessity of drug modification, (6) minimal systemic exposure and (7) avoidance of the liver and gut first-pass metabolism. Neuroprotective cytokines similar in size to LIF (20 kDa), including brain derived neurotrophic factor (14 KDa), ciliary neurotrophic factor (23 kDa), nerve growth factor (13 kDa) vascular endothelial cell growth factor (38 kDa) and heparin-binding epidermal growth factor (22 kDA) have been successfully introduced into the brain via intranasal administration (14). Other studies have demonstrated that IN delivery of insulin-like growth factor-1 and epidermal growth factor promote the proliferation and migration of subventricular zone progenitors in models of demyelination (15, 16). While most of these studies have been performed using rodents, studies have demonstrated the efficacy of intranasal delivery in non-human primates (17, 18) and humans(14, 19, 20).

LIF is the most pleiotropic member of the interleukin-6 family of cytokines, whose expression is induced by injury (21, 22). In the Central Nervous System(CNS), LIF is predominantly released from astrocytes in response to increased neural activity where it promotes oligodendrocyte differentiation and neuronal survival (23). Its expression also increases after CNS injuries that include ischemic stroke, traumatic brain injury, perinatal hypoxia-ischemia, spinal cord injury and in diseases such as multiple sclerosis, Alzheimer’s and Parkinson’s disease (24–29). LIF haplodeficient mice exhibit desynchronized glial responses, increased neurodegeneration, decreased corpus callosum (CC) conductivity and motor deficits after a mTBI (25). LIF has been shown to peak in the acute/subacute phase (at 48 hours) after a mTBI (25), suggesting its possible involvement in the repair and regenerative processes within the injured brain. Given the beneficial role of LIF during the acute/subacute recovery phase after CNS injury and its pleiotropic effects in the brain (30), we hypothesized that IN-LIF treatment would prevent neurodegeneration when administered during the chronic period of recovery from an mTBI. We initiated these studies to test that hypothesis.

## Methods

### Animal Models and ethical approvals

All experiments were performed in accordance with research guidelines set forth by the Institutional Animal Care and Use Committee of Rutgers New Jersey Medical School (Protocol #999900840) and were in accordance with the National Institute of Health Guide for the Care and Use of Laboratory Animals (publication No. 80-23) revised in 1996 and the ARRIVE guidelines.

### mTBI Induction

Male CD1 mice were randomly assigned to sham or injured groups. The mTBI model that we used entails administering a single closed head injury. This model was developed in Dr. Regina Armstrong’s laboratory, as described in Mierzwa et al., 2015 (31). Briefly, 2-month-old mice were administered 0.5 mg/kg Buprenorphine HCL sustained release prophylactically and then anesthetized with 2.5% isoflurane. Once they were confirmed to be deeply anesthetized, the mice were administered Bupivacaine (2.0 µg/g) subcutaneously on the scalp and then a 1 cm midline incision was made to expose the skull. After deflecting the periosteum the animals were placed in a mouse stereotactic device with ear cuffs. The cortical impact device (Custom Design & Fabrication, Richmond, VA), outfitted with a 3-mm-diameter flat metal impactor tip was zeroed on the sagittal suture just in front of Bregma. Twenty seconds after removing the anesthesia, the impactor was driven perpendicularly onto the exposed sagittal suture at a velocity of 4.0m/sec to a depth of 1.3 mm farther than the zero point with a dwell of 100 ms. At the time of impact the mice are immobile and they do not respond to a toe pinch. The mouse was removed from the stereotactic device and the righting time and apnea times were recorded. The animals were re-anesthetized so that the scalp incision could be closed. The mice were then administered 0.9% saline (3% of body weight), i.p. Sham-injured mice were subjected to the same procedures without receiving an impact. All animals were allowed to recover on a heating pad set at 37°C, and upon becoming ambulatory were returned to their home cages. Approximately 8% of the mice in this study died immediately after this TBI.

### IN-LIF administration

Mice were lightly anesthetized with isofluorane so that they would lie on their backs. Mice were administered either 20 µL of recombinant murine LIF (rmLIF) (EMD Millipore, cat #LIF2010) or water using a fine plastic pipette in 2 ul doses. These substances were administered to alternate nostrils within 10 minutes. Two doses of rmLIF were administered, either 40 ng or 100 ng, administered twice daily and 5 days/week for two consecutive weeks during the 6-8 weeks following mTBI.

### Behavioral tests (Irregular horizontal ladder and Catwalk)

Mice were evaluated using the Horizontal Ladder with irregular rungs which we have found to be a very sensitive test of sensorimotor function (32). They were assessed at 4 and 8 weeks post-mTBI. Mice were trained to ambulate across a horizontal ladder with irregularly spaced rungs for 3 days prior to testing **(Fig 1A)**. The task was conducted 3 times non-consecutively per session for each mouse. Video recordings were made and scored by an individual blinded to the group identity of the subject. The mean number of missteps and the average latency to cross the length of the apparatus were determined as the dependent measures for sensorimotor function.

**Figure 1.**
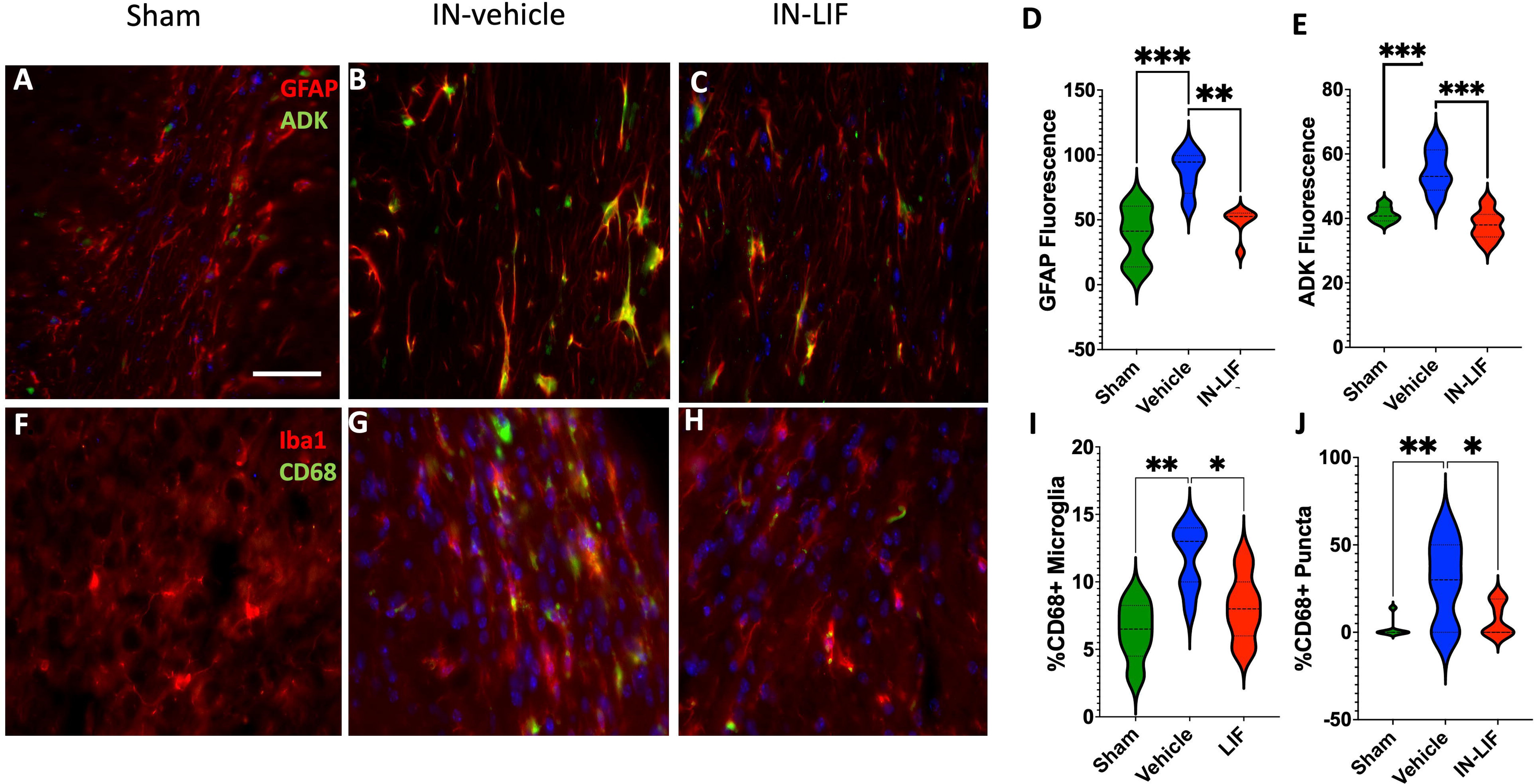
A. IN-LIF Rx improves motor behavior in mTBI mice. The pattern of the irregular horizontal ladder is used to quantify fine motor function. Mice were trained for 3 days on the horizontal ladder before performing the test. **B, C.** Average number of foot slips after 3 non-consecutive runs on the horizontal ladder 4 weeks after the mTBI but before treatment and 8 weeks after the mTBI following 2 weeks of vehicle or 40ng LIF treatments between 6-8 weeks post-TBI. (*p<0.05, n=4-8) by one-way ANOVA followed by Tukey’s multiple comparison test. **D, E.** Catwalk gait analysis measurements showed no differences in mean body speed or regularity index at 8 weeks post-mTBI between groups; one-way ANOVA followed by Tukey’s multiple comparison test, n=4-8 per group.

The mice were also tested using the Catwalk xT. Catwalk xT is a gait analysis software that quantifies gait and footprints in animals that move voluntarily across a glass plate walkway. The animals are video recorded from under the glass plate for each run, or each time the animal walks from one end of the walkway to the opposite end. Each video was classified using the Catwalk software first using the auto classification option, which labeled each foot-print with what the system perceived as being the respective side and position of the animal’s footprints. After the runs were auto classified, they were reviewed to correct errors in the classification and to reduce noise that interrupted appropriate classification. We required three compliant runs for each trial, whch entailed 5-6 full step cycles per run. In our experimental setting we had the minimum duration set to 0.5 s and the maximum duration set to 5 s. Speed variation was set to 60% variation. We used a green intensity value of 0.25. Catwalk xT offers over 200 parameters to assess gait. The parameters of interest focused for this study were maximum contact area, print position, initial dual stance and terminal dual stance.

After testing, the mice were perfused with PBS pH 7.4 containing 4% paraformaldehyde (PFA). The brains were removed and stored in 4% PFA at 4° C overnight and then changed to phosphate buffered saline.

### Magnetic Resonance Acquisition and processing

Two weeks prior to the magnetic resonance imaging (MRI) scans brain samples were washed and stored in fresh PBS containing 0.05% sodium azide, which was changed every week to wash out the fixative to prevent interference with MRI relaxation times (33). One hour before the scans, brains were washed three times for 10 min in 10 mL PBS to remove the PFA solution. Then, the brains were placed into a custom-built MRI-compatible tube filled with Fluorinert, an MRI susceptibility-matching fluid that generates no MRI signal (Sigma-Aldrich, Inc., St. Louis, MO). Microstructural integrity was assessed using ex-vivo diffusion tensor imaging (DTI) of the brain samples using a horizontal bore 9.4T scanner (Bruker BioSpin, Billerica, MA). Proper alignment of each brain sample was performed by keeping the distance from the opening of the tube consistent across all specimens during positioning in the birdcage ^1^H radio frequency coil (15 mm diameter positioning and power optimization were optimized through Bruker defined gradient echo (GE) and fast spin echo (FSE) sequences. Shimming was done by second-order shim with cubic voxel to bring the H20 line width to less than 30Hz. A fast low-angle shot (FLASH 3D) spin echo sequence was used for the T1 images with an FOV of 12×14mm and matrix 120×40, containing 80 slices of thickness of 0.1 mm, TR/TE=50/6 ms. The rapid acquisition with relaxation enhancement (RARE) sequence was used for T2 weighted imaging with a FOV of 12×40, containing 80 slices of thickness of 0.1 mm, TR/TE=50/6 ms. The rapid acquisition with relax8 mm and matrix 102×40, containing 80 slices of thickness of 0.1 mm, TR/TE=50/6 ms. The rapid acquisition with relax64, containing 64 slices of thickness 0.24 mm, TR/TE=3354/24 ms, and number of echoes=8. DTI acquisitions were performed using a Stejskal-Tanner spin-echo diffusion-weighted with a diffusion gradient duration of 4ms and a delay between the two diffusion gradients of 9.3ms. 64 contiguous slices of 240μm thickness were acquired with a field of view of 12 mm×8 mm with a matrix of 102×67, resulting in an in-plane resolution of 120μm × 120μm. TR/TE was 2000/20m with 30 diffusion weighted images and 5 images with no diffusion weighting were acquired for each slice. The diffusion gradient directions for the diffusion weighted images were non-collinear and had the same b-value, 1000 s/mm^2^. Eight averages were acquired with a total experiment time of 10 hours and 24 minutes. After completing the behavioral tests, the mice were perfused with paraformaldehyde, and their brains were removed. They were washed in PBS for 2 weeks to remove the paraformaldehyde before ex-vivo MRI measures.

### Histological staining and analysis

The fixed brains were cryoprotected for 36 h in 30% sucrose in water, frozen in OCT embedding medium on a dry-ice/ethanol slush and then sectioned on a Leica Cryostat at a thickness of 20 µm. Sections were mounted onto SuperfrostPlus slides (VWR, *Radnor,PA*) and stored at −30°. Immunofluorescence was performed as previously described (25). No signal above background was obtained when the primary antibodies were replaced with pre-immune sera. The antibodies used for these studies are listed in Table 1. Images were collected using a Q-imaging Retiga-2000R CCD camera (Surrey, BC, Canada) on an Olympus AX70 microscope (Center Valley, PA, USA). One image from each hemisphere was collected from at least 2 brain sections per mouse. Images were coded so that the analyst was blinded to the specimen’s group identity. Images were analyzed using FiJi software in automated batch mode using a custom script. Briefly, to measure intensity, RGB channels were split, the background was subtracted by applying a rolling ball radius, and the individual channel intensity was read. For % Area and particle count analyses, images were further thresholded, converted to mask, and outlined. Then the “Analyze particles” command was run defining size (0-infinity) and circularity (0-1) that produced a summary file with particle count and % Area values. Images were randomly checked in all 3 groups to ensure the set parameters did not result in false positives or negatives. Images were quantified for ionized calcium-binding adaptor molecule 1 (Iba1) positive microglia and for Iba1/CD68 double positive microglia. Images were manually counted by an analyst blinded to the sample’s group identity. Cells were classified as CD68+ if they had 3 or more bright CD68+ puncta inside of them. White matter astrocytes were classified as adenosine Kinase (ADK)+/Glial fibrillary acidic protein (GFAP)+ if they had a stellate morphology and appeared yellow on the overlay of the green fluorescence for ADK and red fluorescence for GFAP. Images quantified for ADK+/GFAP+ were also coded and manually counted. The scripts used for this manuscript are available at Github: https://github.com/swevison/Levison-Lab-Scripts.

**Table 1.** List of antibodies used for immunofluorescence.

| **Antibody** | **Host** | **Application and dilution** | **Source** | **Catalog #** |
| --- | --- | --- | --- | --- |
| Glial fibrillary acidic protein (GFAP) | Rabbit | Immunofluorescence, 1:200;<br>Western blot, 1:1000 | Dako, Santa Clara, CA | Z0334 |
| Amyloid precursor protein (APP) Y188 | Rabbit | Immunofluorescence, 1:200 | Abcam, Cambridge, MA | ab32136 |
| Ionized calcium binding adaptor molecule 1 (IBA1) | Rabbit | Immunofluorescence, 1:200 | Wako Pure Chemical Corporation, Wako, TX | 019-19741 |
| CD-68 | Rat | Immunofluorescence, 1:200;<br>Western blot, 1:1000 | Bio-Rad, San Diego, CA | MCA1957GA |
| Adenosine Kinase (ADK) | Rabbit | Immunofluorescence, 1:500 | Fisher Scientific, Pittsburgh PA | 501567320 |
| RedX-conjugated anti-mouse IgG | Donkey | Immunofluorescence, 1:300 | Jackson ImmunoResearch, West Grove, PA | 715-295-150 |
| Alexa 488-conjugated anti-rabbit IgG | Donkey | Immunofluorescence, 1:300 | Jackson ImmunoResearch, West Grove, PA | 711-545-152 |
| Alexa 594-conjugated anti-rabbit | Donkey | Immunofluorescence, 1:300 | Jackson ImmunoResearch, West Grove, PA | 711-585-152 |
| Alexa 488-conjugated donkey anti-mouse IgG | Donkey | Immunofluorescence, 1:300 | Jackson ImmunoResearch, West Grove, PA | 715-545-150 |

### Statistical analyses

All sham T2 weighted images were registered to the mouse Franklin and Paxinos MRI atlas and averaged to obtain a study anatomical template (34). Subsequently all anatomical T1, T2 and DTI images were spatially registered to the anatomical study brain template using non-linear registration in Bioimage Suite (35). The resulting brain matrix volume data were used to calculate the DTI eigenvalues (λ_1_, λ_2_, λ_3_) and vectors (e_1_, e_2_, e_3_). At each voxel, the eigenvalues represented the magnitude of diffusion and eigenvectors represented maximal and minimal diffusion directions respectively. The fractional anisotropy (FA) and mean diffusion (MD) were estimated from the eigen values (36, 37) using the BioImage Suite software (35).

Statistical parametric mapping of the whole brain was performed in a voxel-wise manner using a two sample t-test (two-tailed) using AFNI (38). Statistical differences between experimental groups were analyzed with a corrected threshold probability of P<0.05, determining voxels with a significant difference (active voxels) using a family wise error control of a minimum cluster of 30 contiguous voxels to correct for multiple comparisons. MRI intensity differences obtained from the t-test means were depicted in color, with hot colors indicating a positive mean difference, and cold colors depicting a negative mean difference, which were overlayed on the mean anatomical MRI of the control group animals. Regions of interest (ROI) analysis was manually performed by drawing regions using the Franklin Paxinos mouse stereotactic atlas (34). Average DTI parameters across each ROI were determined across each animal subject in the respective experimental groups. Group-level statistical comparisons were performed using a one-way ANOVA followed by a post-hoc Tukey’s HSD test with a significance of p<0.05 required for significance.

Raw data from image analyses and behavioral tests were imported into Prism (GraphPad 10 Software; La Jolla, CA) for statistical analyses using One-way ANOVA followed by Tukey’s post hoc intergroup comparison. Graphs were produced in Prism and error bars denote standard error of means (SEMs). Comparisons were interpreted as significant when associated with p<0.05.

## Results

### Delayed LIF treatment prevents sensorimotor behavior debilitation after mTBI

Horizontal ladder tests were performed at 4 and 8 weeks post-mTBI in naïve, sham, and mTBI mice with intranasal-vehicle (IN-vehicle) or IN-LIF. Between 4 weeks and 8 weeks after a mTBI the motor performance of the mice that were administered the vehicle showed significant functional deterioration on the horizontal ladder as evidenced by increased numbers of foot slips (compare **Fig 1B** and **1C**; blue). While only 10% of animals showed an average of >2 foot slips at 4 weeks post-mTBI **(Fig 1B**; blue), 50% of animals showed >2 foot slips at 8 weeks post-injury **(Fig 1C**; blue), indicating delayed sensorimotor deterioration. Remarkably, all of the mTBI mice treated with IN-LIF showed <2 foot slips at 8 weeks post-injury, a level of motor performance that was no worse than the sham injured mice and significantly better (p < 0.01) than the vehicle administered mTBI mice **(Fig 1C**; red). Catwalk gait analysis showed no differences in any of the evaluated parameters (**Fig 1E-F**). For example, there was no difference in mean body speed or regularity index between groups at 8 weeks post-injury **(Fig 1E** and **1F)**. Interestingly, the female mice showed spontaneous recovery at 8 weeks on the irregular horizontal ladder and did not benefit from IN-LIF Rx (data not shown). Therefore, only young adult male mice were used for the remainder of these studies.

While the majority of people who experience a concussion will have just one in their lifetime, a smaller proportion of individuals will sustain recurrent concussions, which represents another clinically important subset. Therefore, initiated studies using a repeated mild TBI (rmTBI) model using the parameters described by Hall et al., 2016 (39) to enable us to evaluate the therapeutic benefit of delayed IN-LIF treatment. Using this rmTBI model we established that the mice subjected to a rmTBI had impaired motor performance 3 days after the last concussive injury (**Fig S1**) and that these mice had both increased astrogliosis in both the gray matter and white matter as determined by increased GFAP **(Fig S2)** and decreased adenosine kinase **(Fig S3)**, along with increased microgliosis as determined by increased percentage of CD68+/Iba1+ cells in the injured neocortex (**Fig S4**). However, when their motor performance was evaluated over 6 weeks of recovery, they showed almost full recovery (**Fig S5**). Since the goal of the current studies was to evaluate the neurorestorative properties of IN-LIF during the chronic period of recovery from mTBI, we did not pursue studies using the rmTBI model further.

### Evidence of LIF neuroprotection from T1 and T2 contrast MRI changes after mTBI

Conventional MRI T1 and T2 weighted acquisitions were obtained using the FLASH and RARE sequences, respectively **(Fig S6)**. Comparing the T1 contrast changes between the mTBI+vehicle treated versus sham group, significant clusters of T1 decrease were observed across the neocortex proximal to the TBI epicenter along with diffuse clusters of T1 decreases within the striatum, basal brain areas and the corpus callosum **(Fig 2A)**. Relatively fewer T2 intensity changes were detected after mTBI, with T2 decreases observed in the neocortex adjacent to the mTBI epicenter compared to sham **(Fig 2D)**. IN-LIF treatment significantly reduced the mTBI-induced T1 intensity decreases across most frontal regions of the brain and adjacent to the mTBI epicenter, in addition to increasing the T1 intensity across the basal forebrain, somatosensory and entorhinal cortices **(Fig 2B)**. T2 intensity decreases observed after mTBI were absent with IN-LIF treatment, and increased T2 was observed across the frontal corpus callosum **(Fig 2E)**. The areas affected by IN-LIF were determined by comparing mTBI groups treated with vehicle versus LIF. As shown in **Fig 2C**, the basal forebrain, somatosensory cortex, and lateral ventricles showed significantly reduced T1 intensity in mTBI+vehicle treated mice compared to mTBI+IN-LIF mice. T2 was relatively less sensitive than T1, and the IN-LIF effects were only evident in the lateral ventricular space in the mTBI+IN-LIF treated vs. the mTBI+IN-vehicle treated mice **(Fig 2F)**. Sparse T2 decreases were observed across the corpus callosum in the mTBI+IN-vehicle mice compared to mTBI+IN-LIF mice **(Fig 2F)**, signifying the protective effect of IN-LIF over the corpus callosal area.

**Fig 2.**
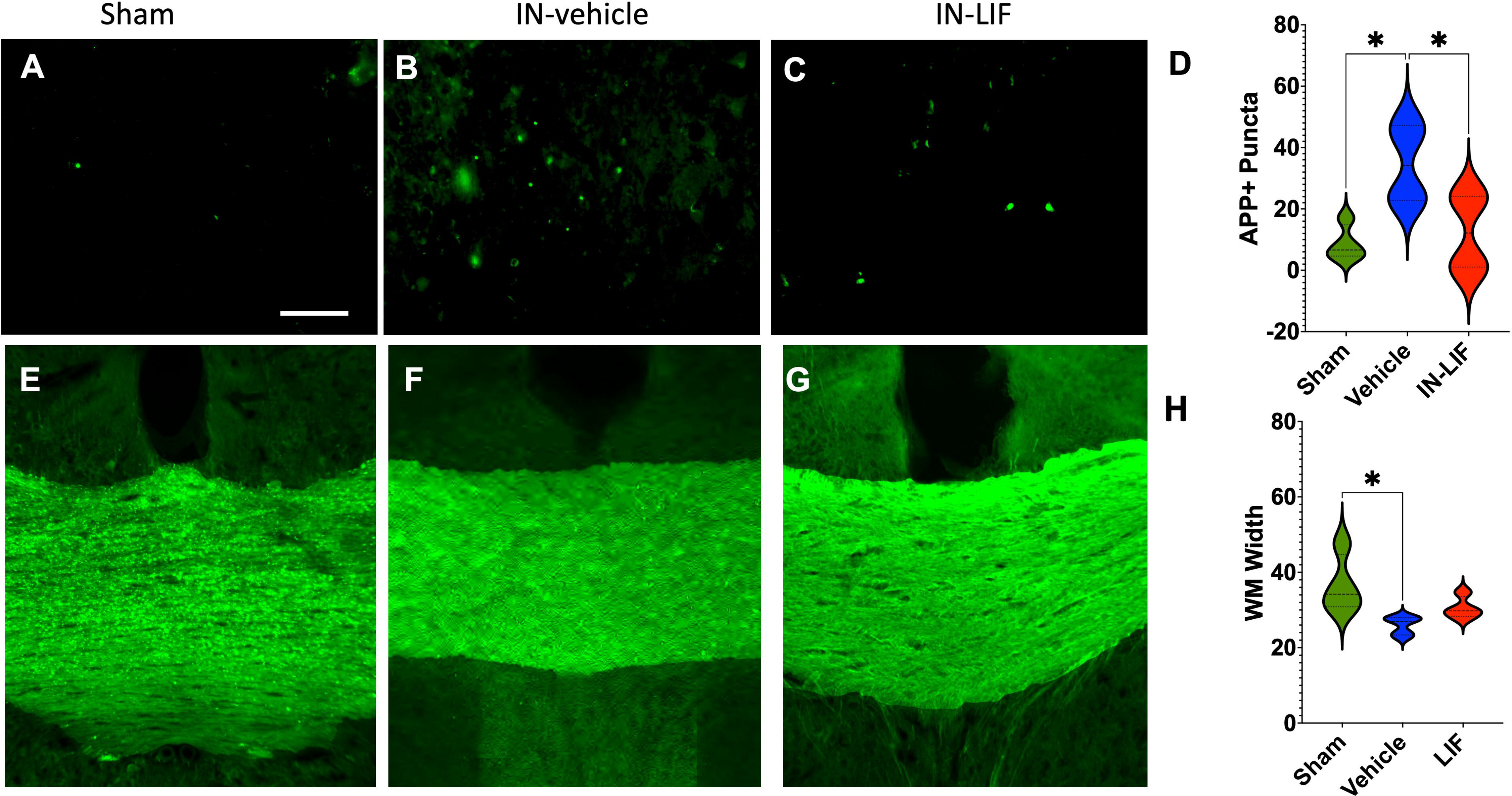
IN-LIF Rx modifies the mTBI-induced MRI-T1 and T2 intensity changes. Group differences in T1 (FLASH) and T2 (RARE) signal intensities were estimated using a two-sample t-test on a voxel-by-voxel basis across the whole brain. A,D mTBI mice treated with vehicle vs sham mice, B,E. LIF-treated mTBI mice vs sham mice and C,F. vehicle-treated mTBI mice vs LIF-treated mTBI mice. Activated voxels have a threshold of P<0.05, corrected for multiple comparisons using a family-wise error control of 20-voxel contiguous cluster volume. Hot colors depict larger T1 or T2 intensity across the first group vs the second group. Sham mice (n=7), mTBI mice treated with vehicle (n=7) and mTBI mice treated with LIF (n=8). Red arrows demark the mTBI epicenter.

### IN-LIF partially restores callosal DTI changes after mTBI

Following the conventional T1 and T2 acquisitions, diffusion-sensitive acquisition was performed, and DTI parameters such as the fractional anisotropy (FA) and mean diffusivity (MD) were estimated across the whole brain **(Fig 3)**. Comparing group differences between mTBI mice treated with vehicle and sham mice, FA decreased predominantly in the corpus callosum and other subcortical white matter areas **(Fig 3A**; vehicle vs sham). MD increased between vehicle-treated mTBI mice and sham mice and was prominent across the neocortex adjoining the impact epicenter and deeper areas such as the corpus callosum, subventricular zone and the striatum **(Fig 3B**; vehicle vs sham). IN-LIF treatment inhibited the mTBI-induced FA decrease in a dose-dependent manner **(Fig 3A**; IN-LIF-40 vs sham and IN-LIF-100 vs sham). However, sparse increases in FA were observed in certain deeper areas of the brain such as the internal capsule (IC) and anterior commissure pars posterior (ACP), possibly owing to the proximity of the intranasal pathways and higher LIF dose **(Fig 3A**; IN-LIF-100 vs sham). IN-LIF treatment inhibited the mTBI-induced MD increases in a dose-dependent manner **(Fig 3B**; IN-LIF-40 vs. sham and IN-LIF-100 vs. sham). However, the higher IN-LIF dose increased MD in the subventricular zone and deeper brain regions **(Fig 3B**; IN-LIF-100 vs sham). The T1 decreases over the ventricles as described above overlapped well with the ventricular MD changes that were particularly seen at the epicenter and adjoining coronal slices as seen in **figs 2A and 3B**. These changes were abrogated by IN-LIF (**Figs. 2C and 3B**).

**Figure 3.**
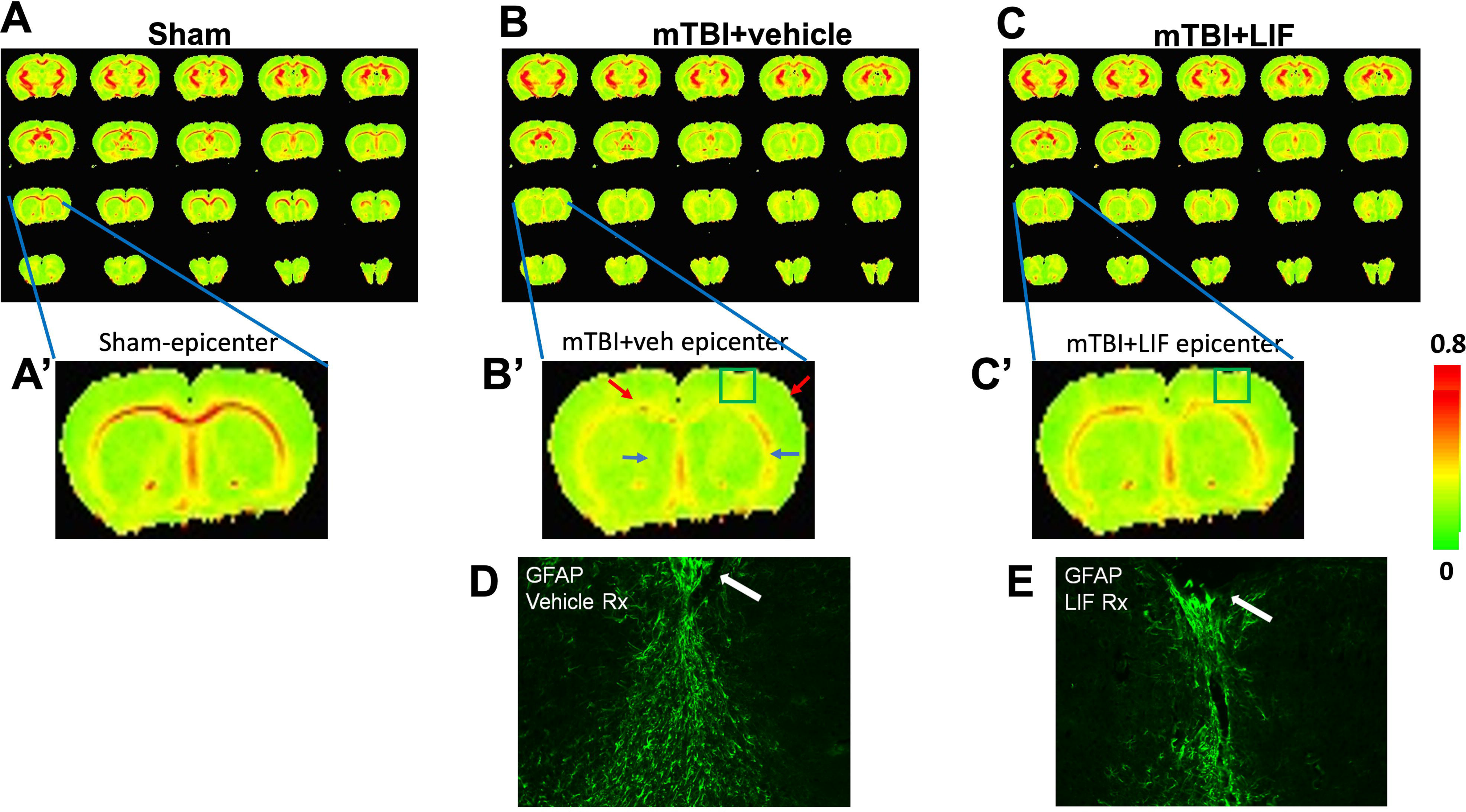
IN-LIF Rx reduces mTBI-induced DTI-FA and MD changes, indicating axonal protection—group differences between sham, vehicle-treated, and LIF-treated mTBI mice. **A.** fractional anisotropy (FA) changes **B.** mean diffusivity (MD) changes. The TBI epicenter is located at +0.5 mm AP from the bregma, as indicated by red arrows. Activated voxels represent a threshold of p<0.05 after a 2-tailed t-test corrected for multiple comparisons using a contiguous cluster volume of 30 voxels. The color map indicates the magnitude of the change in FA or MD. Red arrows demark the mTBI epicenter.

A region of interest (ROI) analysis involving 8 ROIs was performed over the different animal groups. Significant FA decreases were observed across the corpus callosum (CC) and prefrontal cortex (PFC) in the mTBI vehicle group (**Fig 4A,B**, p < 0.001). Significant increases in MD also were observed in the CC and the PFC (**Fig 4C,D**, p < 0.01). No significant changes in FA or MD occurred across 6 other ROIs tested (**Figs S6** and **Fig S7**). However, an increasing trend in MD was observed in the motor cortex, primary somatosensory cortex, striatum, hippocampus, insula, and thalamus **(Fig S7)**. IN-LIF treatment, in a dose-dependent manner, reversed the mTBI-induced FA decrease across the corpus callosum and the PFC, respectively **(Fig 4A,B**, p < 0.001). However, IN-LIF at the lower dose of 40 ng restored the mTBI-induced MD increase observed across the corpus callosum and in the PFC, with no further improvements with the 100 ng dose **(Fig 4C,D**, p < 0.01). When FA intensity was plotted against the number of foot slips on the horizontal ladder for Vehicle Rx TBI mice vs. LIF Rx TBI mice a Pearson’s correlation analysis showed that there was a small but significant correlation for the LIF Rx mice FA and numbers of footslips (p<0.05, R2 = 0.53), whereas there was no correlation between the FA for the Vehicle Rx mice. The elevations were statistically different (F=17.4; p =0.001) (**Fig4E**). We also plotted the FA intensity against the number of APP+ puncta evident by immunofluorescence staining, as depicted in Fig. 6, using brain sections from Vehicle Rx TBI mice and LIF Rx TBI mice. A Pearson’s correlation analysis showed that there was a correlation between the FA and the APP+ puncta for theVehicle Rx mice (p < 0.05, R2=0.66) whereas there was no correlation between the FA for the LIF Rx mice. Again, there was a significant difference in the elevation (F=22.8, p=0.0005) **(Fig4F)**. When FA intensity was plotted against GFAP fluorescence intensity, as depicted in Fig. 7, a Pearson’s correlation analysis showed that there was a significant correlation for the LIF Rx TBI mice vs. GFAP+ intensity (P<0.01, R2= 0.77) but no correlation for the Vehicle Rx TBI mice. Again, the elevations were statistically different (F=25.3; p <0.0005).

**Figure 4.**
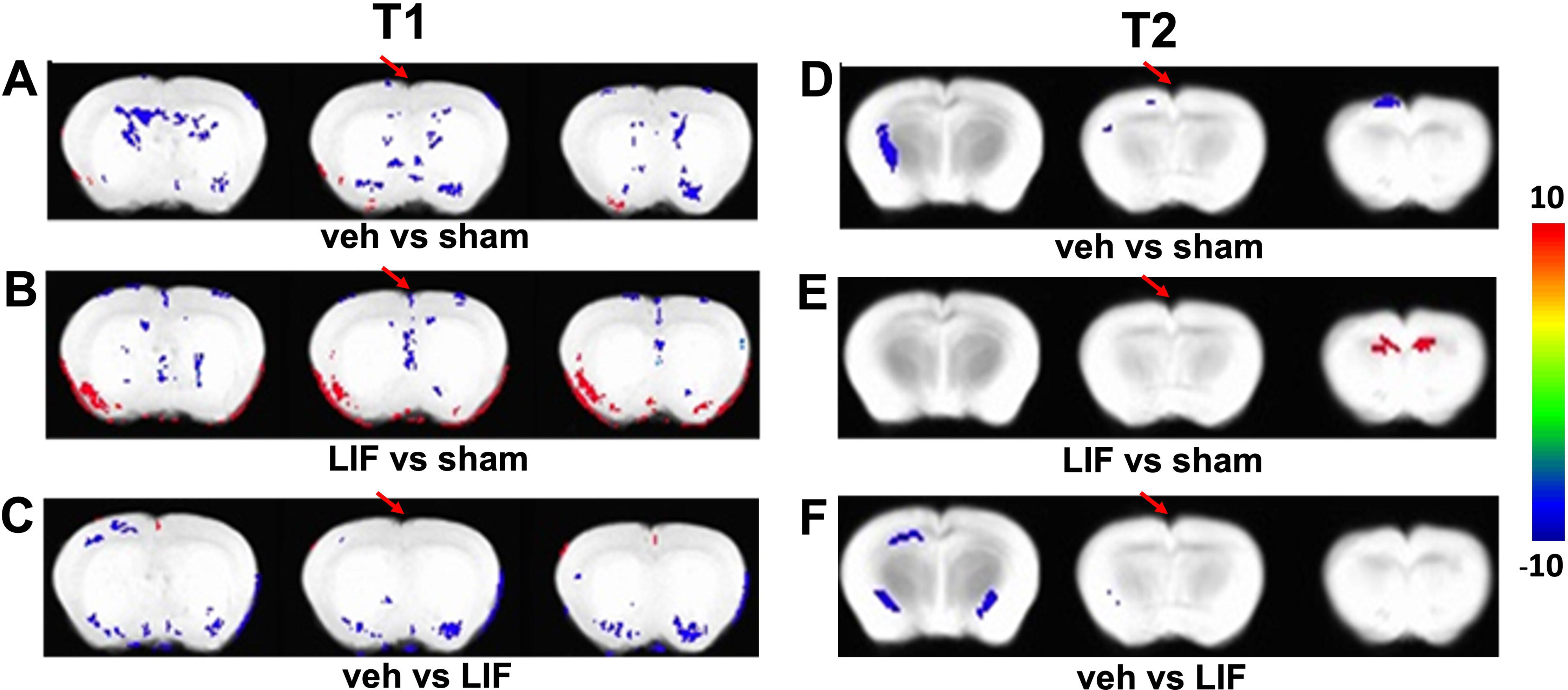
IN-LIF Rx reduces mTBI-induced DTI-FA changes in a dose-dependent manner. Quantitative DTI parameter changes across the corpus callosum (CC) and prefrontal cortex (PFC) regions of interest in sham (blue; n=7), mTBI+vehicle (orange; n=8), mTBI+40ng LIF (gray; n=8) and mTBI+100ng LIF (yellow; n=9). **A,B.** FA and **C,D.** MD. One-way ANOVA followed by a post-hoc Tukey’s HSD test with p<0.05 was required for significance between groups. **A.** F=13.63, **B.** F=12.8, **C.** F=6.74 and **D.** F=5.47. **E.** FA intensity was plotted against the number of foot slips on the horizontal ladder for Vehicle Rx TBI mice vs. LIF Rx TBI mice. A Pearson’s correlation analysis showed that there was a small but significant correlation for the LIF Rx mice FA and numbers of footslips (p<0.05, R2 = 0.53), whereas there was no correlation between the FA for the Vehicle Rx mice. The elevations were statistically different (F=17.4; p =0.001). **F.** FA intensity was plotted against the number of APP+ puncta for Vehicle Rx TBI mice and LIF Rx TBI mice. A Pearson’s correlation analysis showed a correlation between FA and APP+ puncta for theVehicle Rx mice (p < 0.05, R2=0.66) but no correlation between the FA for the LIF Rx mice. There was a significant difference in the elevation (F=22.8, p=0.0005). **G.** FA intensity was plotted against GFAP fluorescence intensity. A Pearson’s correlation analysis showed a significant correlation for the LIF Rx TBI mice vs. GFAP+ intensity (P<0.01, R2= 0.77) but no correlation for the Vehicle Rx TBI mice. The elevations were statistically different (F=25.3; p <0.0005).

Compared to the sham mice, TBI mice showed decreased FA across the corpus callosum and other white matter tracts and increased FA across neocortical and subcortical gray matter regions **(Fig 5A,B)**. When the mTBI mice were treated with 40 ng IN-LIF, the FA decrease across the corpus callosum was inhibited and the neocortical and subcortical gray matter FA increases were also prevented **(Fig 5C)**. To ascertain the underlying cellular changes representing these FA increases in the neocortex, the brains after the ex-vivo DTI studies were stained for GFAP as an index of astroglial reactivity. Astroglial scars were evident at the dorsal edges of the site of impact from the mTBI, which was mitigated by 40 ng IN-LIF treatment **(Fig 5D)**, linking glial reactivity to the observed neocortical FA increases around the mTBI epicenter and subcortical gray matter areas.

**Figure 5.**
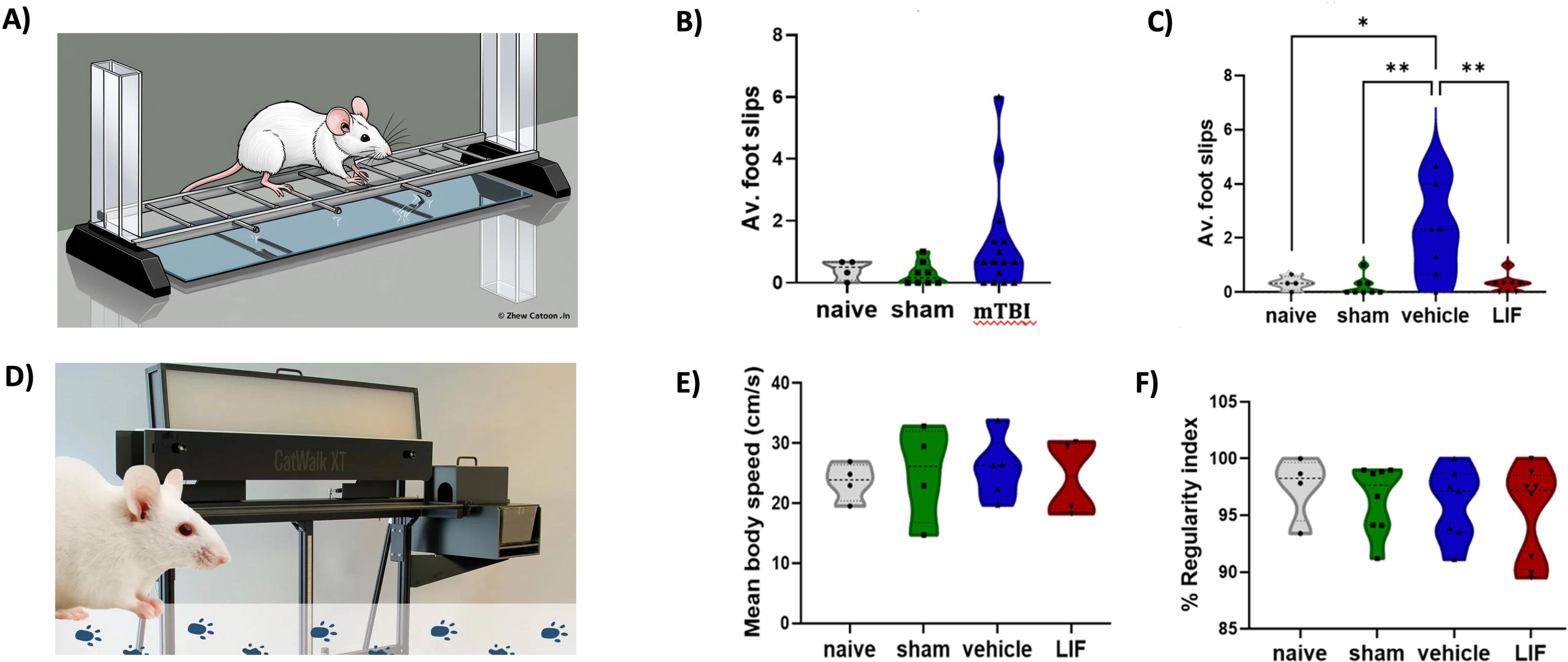
DTI-FA captures glial reactivity after mTBI, which IN-LIF Rx mitigates. Pseudocolor images of the FA maps from **A.** sham, **B.** mTBI+vehicle and **C.** mTBI+LIF groups. Colors represent FA from group averages, sham (n=7), mTBI+vehicle (n=8) and mTBI+LIF (n=8). Increased FA was observed across cortical (red arrows) and subcortical gray matter areas (blue arrows) in the mTBI mice treated with vehicle. **D.** Representative 10X images showing astroglial scar at the impact epicenter from vehicle Rx and LIF Rx mice (white arrows).

### IN-LIF protects against delayed axonal degeneration following mTBI

Axonal transport is compromised in damaged axons leading to the accumulation of amyloid precursor protein (APP) in axonal swellings along the length of the axon. Additionally, axons can be transected during TBI, producing end-bulbs (40). Therefore, we evaluated the extent of axonal damage following mTBI by immunostaining coronal sections using the Y188 antibody that is highly specific for APP (41). Negligible APP staining was seen in the CC in the sham group **(Fig 6A)**. By contrast, APP+ puncta accumulated within CC fibers of the IN-vehicle group as distinct swellings or end bulbs **(Fig 6B)**. 40 ng IN-LIF Rx decreased the number of APP swellings and endbulbs **(Fig 6C)**. A particle analysis showed significantly increased numbers of APP+ particles in the CC of the vehicle group over sham, which was almost entirely reversed by IN-LIF **(Fig 6D**, *p<0.05). To determine whether IN-LIF would also improve the myelin integrity, sections were stained for MBP and the thickness of the corpus callosum was measured (**Fig 6E-H**). This analysis revealed a 30% decrease in the CC thickness in the TBI-vehicle group vs. the sham group (p <0.001) with a 17% decrease for the IN-LIF Rx mice that was not statistically different from the sham group **(Fig 6H)**.

**Fig 6.**
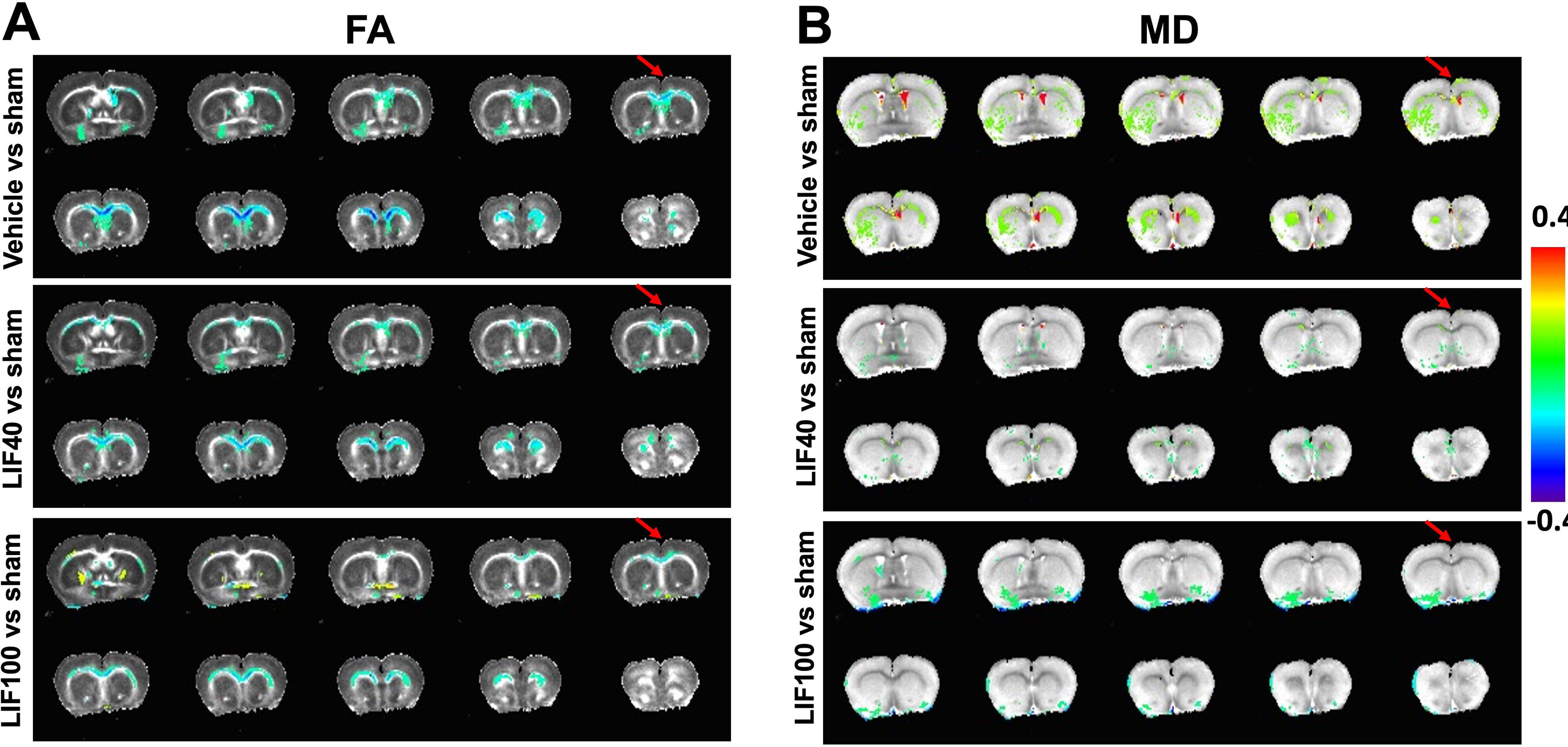
IN-LIF Rx decreases delayed axonal injury in the corpus callosum after mTBI. Mice were injured at P60 and received IN-LIF for 2 weeks as described in Fig 2. Images of APP staining in the CC for sham **(A)**, IN-vehicle treated **(B)** and IN-LIF Rx **(C)** mice. **D) The g**raph depicts particle analysis of APP staining. Sections from sham **(E)**, IN-vehicle treated **(F),** and IN-LIF Rx **(G)** mice were stained for MBP and the thickness of the corpus callosum was measured **(H)**. Scale bar represents 50 µm for panels A-C and 200 µm for panels E-G. Data were analyzed by one-way ANOVA followed by Tukey’s multiple comparison test. Statistical significance is indicated as follows: *p* < 0.05 (*), *p* < 0.01. n=6-8/group.

### Delayed IN-LIF attenuates astrocyte activation in the injured subcortical WM after mTBI

GFAP has long been used as a marker for reactive gliosis, and GFAP expression was significantly increased in the subcortical WM of the vehicle treated group (**Fig 7B**,**D**, p < 0.001), but GFAP is a structural protein that does not provide information about the functional state of the astrocytes. Studies using the controlled cortical impact model have shown that TBI causes the overexpression of ADK in reactive astrocytes, a major adenosine metabolizing enzyme. Increases in ADK will result in chronic adenosine deficiency which can decrease the threshold for seizures (42). Therefore, we stained sections for ADK and evaluated the effect of 40 ng IN-LIF Rx on its expression. The subcortical white matter of the sham group showed low ADK immunofluorescence **(Figs 7A,E**. By contrast, ADK was strongly expressed in the cytoplasm of hypertrophic GFAP+ astrocytes in the vehicle Rx mTBI group **(Fig 7B,E)**. IN-LIF decreased levels of ADK to that seen in the sham injured group which was significantly reduced from those seen in the vehicle-treated group **(Fig 7C,E**, p < 0.01). Not only did IN-LIF Rx decrease the overall level of ADK, but it reduced the percentage of astrocytes that were ADK+ to that seen in the sham-injured mice **(Fig 7I**, p < 0.01 vs. vehicle).

**Fig 7.**
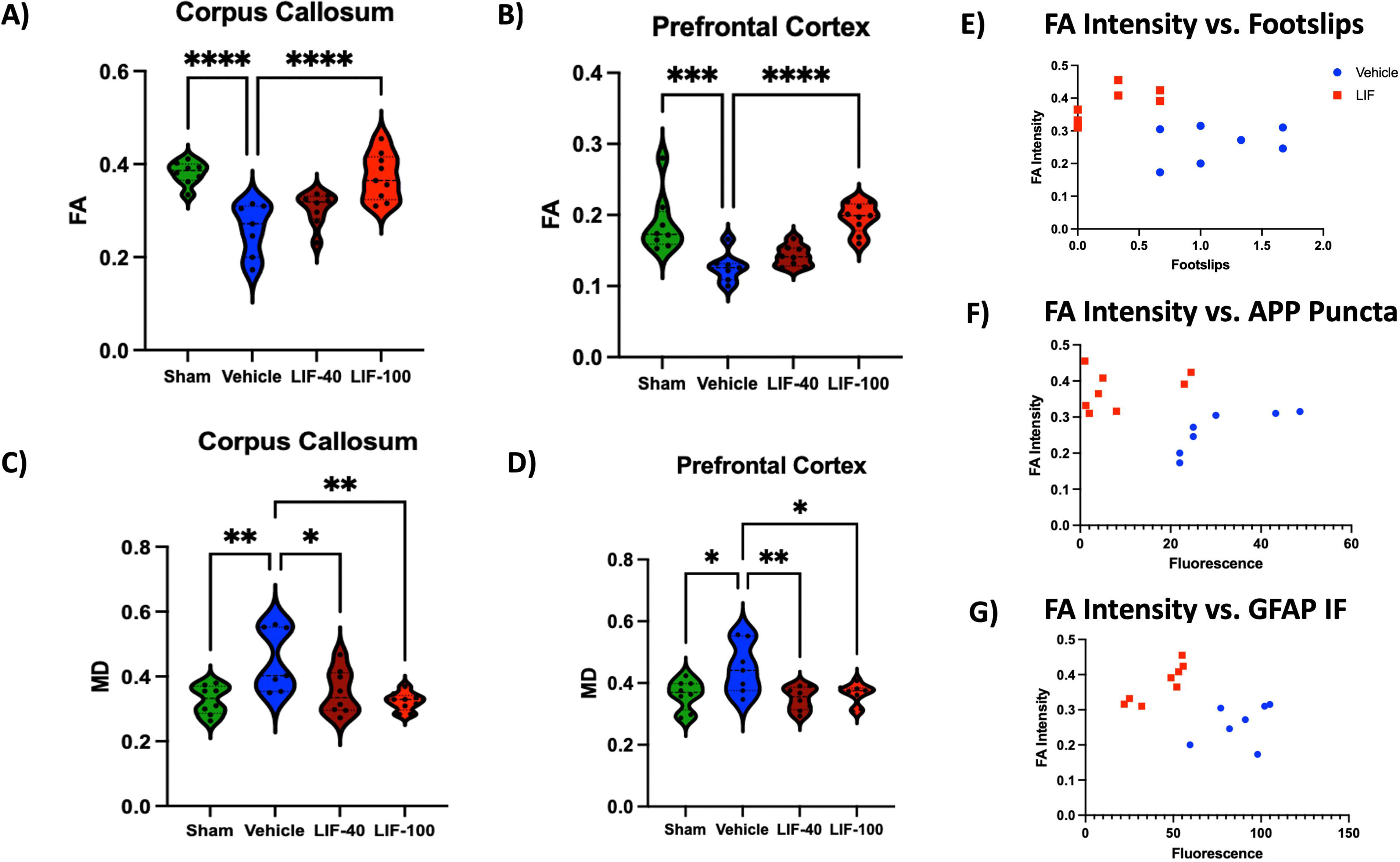
IN-LIF Rx decreases astroglial and microglial reactivity after mTBI. Mice were injured at P60 and received IN LIF for 2 weeks as described in Fig 2. After behavioral analyses, the mice were perfused, the brains sectioned and stained for GFAP, ADK, IBA-1 and CD-68. Scale bar represents 100 µm. Data were analyzed by one-way ANOVA followed by Tukey’s multiple comparison test for GFAP immunofluorescence (D) ADK immunofluorescence (E), % CD68/IBA-1+ (I0 and for CD68 immunofluorescence (J). * p < 0.5, ** p<0.01, ** p<0.005, *** p < 0.001. n=6-8/group.

### Delayed IN-LIF attenuates macrophage/microglial activation in the injured white matter after mTBI

In our studies of mTBI in LIF heterozygous juvenile mice, we observed delayed and amplified microgliosis after injury (25). Therefore, we hypothesized that IN-LIF would repress microgliosis. To test this hypothesis, we immunostained coronal sections for Iba1 and CD68. The sham injured white matter was populated by ramified IBA1+ cells with few CD68+ puncta **(Fig 7F)**. As expected, the subcortical WM of the vehicle-treated mTBI group was populated by IBA1+ cells that also contained multiple lysosomes that labeled for CD68 **(Fig 7G)**. By contrast, most of the microglia in the 40 ng IN-LIF Rx brains had a ramified morphology and they contained few CD68+ lysosomes **(Fig 7H)**. When we quantified the percentage of CD68+/IBA1+ cells in the subcortical WM, on average 30 ± 9.7% of the microglia were double positive in the vehicle-treated mTBI group. In contrast, less than 2.3 ± 2.6% were double positive in the sham injured group and there were 7.8 ± 3.6% double positive in the IN-LIF Rx group (p<0.05 comparing IN-vehicle vs IN-LIF) **(Fig 7J)**.

## Discussion

Histopathological analyses after mTBI have revealed both acute as well as chronic phases of neurodegeneration. To date the majority of pre-clinical studies have focused on strategies to reduce acute brain injury (9) with a paucity of studies evaluating the mechanisms of delayed neurodegeneration and its treatment. In the single hit model that we have employed, the subcortical white matter appears normal for up to 6 weeks after the injury. Still, tertiary neurodegeneration becomes apparent by 8 weeks of recovery. Accompanying the Wallerian degeneration that ensues the compound action potential becomes weak and electron microscopic analyses support the conclusion that the axons are undergoing pathological changes that precede any changes in myelin maintenance (12). Pediatric, juvenile and adult TBI studies have all revealed delayed white matter injury. There are several hypotheses for this late-onset neurodegeneration. One hypothesis is that the white matter becomes hypoxic which triggers the delayed white matter loss (43). Alternatively, the Wallerian degeneration could result from SARM1 activation as delayed axonal degeneration does not occur in SARM1 null mice after either mTBI or repeated head injury (8, 13, 44). While the trigger for the delayed neurodegeneration seen in this model remains elusive, our studies have reproduced the delayed white matter neurodegeneration that is seen after single hit concussive injury as revealed by high-resolution DTI, histopathology for axonal swellings and progressive sensorimotor deficits. Notably, all of these aspects of tertiary neurodegeneration and the accompanying astrogliosis and microgliosis subsequent to the mTBI could be largely prevented (or reversed) by administering LIF intranasally twice daily starting at 6 weeks of recovery.

IN-LIF broadly affected the injured brain, and its effects were especially prominent at the impact epicenter and in specific deeper brain structures as visualized by DTI. Moreover, IN-LIF treatment inhibited the mTBI-induced FA decrease, the MD increase, and there was a strong trend for it to reduce the sensorimotor deficits in a dose-dependent manner. While the IN-LIF had a widespread neuroprotective effect, it appeared to more strongly affect basal brain areas **(Fig 2B)**, likely because those regions have more robust structural connections with the olfactory bulb, which may have resulted in greater bioavailability of IN-LIF in these areas, especially with exposure to the higher dose of IN-LIF. In our studies of pediatric TBI we tested doses of 10 and 20 ng, with the 20ng dose showing statistically significant benefits(32). We calculated that if we administered 40 ng of LIF and if 2% of that LIF entered the brain(45), then given the extracelllar volume of ∼ 100 µL for an adult mouse brain that this would achieve a concentration of 8 ng/mL, which would achieve ∼25% occupancy of the LIF receptor that has a kD of 1.1 nM. Our data show that this dose was clearly neuroprotective; however, we suspected that a higher dose might be more effective; therefore, we increased the dose to 100 ng and found that this provided superior neuroprotection **(Fig 3B)**. Further studies will be required to ascertain whether this is the optimal dose for administering native LIF. However, we have shifted our future studies to using LIF encapsulated into nanoparticles, which will likely have better efficacy with less frequent dosing than native LIF.

A novel finding in the present study was that FA and MD changed across the prefrontal cortex 8 weeks after an mTBI. Furthermore, MD showed a trend for increasing across several gray matter brain regions, indicating milder effects across a large area of gray matter. This increasing trend in MD was significantly reversed by IN-LIF treatment across all of these regions. The prefrontal cortex plays a vital role in social cognition in humans and across animal models (46). Changes in social cognition have been observed after TBIs (47) stemming from post-injury frontal lobe changes and prefrontal cortical reorganization (48). While our behavioral paradigms did not test for differences in social cognition after mTBI, the current DTI findings, indicating a significant impact on the prefrontal cortex, warrant behavioral tests of social cognition during the delayed phase of recovery from a mTBI.

An additional finding was the formation of glial scars across the cortical gray matter adjoining the mTBI epicenter (ascertained by GFAP staining; **Fig 5D**), which manifested as areas of increased FA at the margins of the TBI epicenter **(Fig 5B**; arrows). The mTBI-induced glial scar formation was prevented after treatment with 40 ng IN-LIF **(Fig 5C)**. Within the white matter, the IN-LIF Rx mice also showed reduced astrogliosis as illustrated by fewer GFAP+/ADK+ stellate astrocytes. As ADK overexpression will reduce levels of adenosine, which lowers the threshold for epileptic seizures through insufficient activation of adenosine A_1_ receptors (42), our data show that not only does IN-LIF reduce the formation of the glial scar, but it also enhances a key function of the astrocytes that should reduce seizure incidence.

Microglia become activated rapidly after brain injuries and produce chemokines that attract macrophages, contributing to secondary and tertiary brain injury (49, 50). Therefore, we assessed the functional state of the microglia/macrophages using antibodies to Iba-1 and CD-68. CD-68 is a lysosomal protein that participates in antigen processing in late endosomes and the degradation of proteins in lysosomes. Therefore, it is a useful marker for highly phagocytic, activated microglia. As the microglia were significantly less reactive using delayed 40 ng IN-LIF treatment, a reasonable conclusion is that the production and release of cytokines and chemokines from the microglia have been suppressed and, therefore, they are no longer stoking an inflammatory response. Indeed, if LIF has shifted the microglia towards an M2-like phenotype, then they are likely releasing growth factors and trophic factors that are facilitating repair. Reminiscent of our results, systemically administered LIF reduces the numbers of activated macrophage/microglia in the ischemic rat brain and LIF has been shown to reduce macrophage IL-12 levels in vitro (51).

## Conclusions

Using T1/T2 MRI and DTI imaging we have obtained clinically relevant markers of gray matter and white matter changes across the brain in a mouse model of mTBI. The deterioration of sensorimotor function seen between 6 and 8 weeks of recovery correlated with white matter loss across the corpus callosum and in the PFC. Combining behavioral analyses, histology and translational imaging, our studies document the restorative effects of intranasally administered LIF. The dose-dependent effects of LIF in improving white matter integrity suggest that strategies that can increase LIF bioavailability (e.g., nanoparticle encapsulation) may improve its translational and clinical trial prospects as a treatment for preventing tertiary neurodegeneration.

### Transparency, Rigor and Reproducibility Summary

Sample size 6-8 mice per group, were based on our preliminary studies with observed differences between means (>1.5 fold), observed standard deviations (40% of lowest mean) which indicated >80% power to detect both an overall significant effect by ANOVA and a post-hoc difference between the experimental groups vs. control with p<0.05 after correction for multiple comparisons. Thirty mice were subjected to the single hit mTBI model, and 14 mice were subjected to the rmTBI protocol. None were excluded for technical reasons, and 2 mice died immediately after the mTBI, and one mouse died after the rmTBI. TBI mice were randomly assigned to IN-vehicle or IN-LIF Rx groups. The investigators who performed the behavioral tests were blinded to the group identity, as were those performing the MRI analyses. The normality of the behavioral and histological outcome data were evaluated using Shapiro-Wilk tests. The preprocessed MRI and DTI data sets cannot be shared in public due to ongoing therapeutic development of LIF. Still, data will be made available to readers upon receipt of a request by the corresponding author.

## Supporting information

D'Mello Fig. S1

D'Mello Fig. S2

D'Mello Fig. S3

D'Mello Fig. S4

D'Mello Fig. S5

D'Mello Fig. S6

D'Mello Fig. S7

D'Mello Fig. S8

## Contributors

Conceptualization: SWL, SSK and FH; funding acquisition: SWL, SSK and FH; project supervision: SWL, SSK and FH; experimental data acquisition: VD, AM, SA, MF, JM, BGS, DC and YW; behavioral data analysis: VD, AM and SWL; Immunofluorescence analyses: VD, MF and SWL; MR image processing and analysis: JM, BGS and SSK; writing original manuscript draft: VD, SWL and SSK. Reviewing and editing revised versions: SWL, SSK and FH.

## Supplemental figures and legends for D’Mello et al., 2025

**Supplementary Figure S1**. rmTBI mice take significantly more time than the sham to exit an enclosure. A) Quantification of the time that each mouse took to exit the circle. n=4/group, *p<0.05 by Student’s t-test. B) A schematic of the exit circle apparatus.

**Supplementary Figure 2.** Activated astrocytes are increased at 72 h after rmTBI. Immunofluorescence for GFAP (red) in the cerebral cortex (A,B) and the corpus callosum (C,D). Data were analyzed by one-way ANOVA followed by Tukey’s multiple comparison test. Statistical significance is indicated as follows: p < 0.05 (*), p < 0.01 (), p < 0.001 (***) n=4/group. Scale bar represents 50 µm.

**Supplementary Figure 3.** Adenosine kinase (ADK) is reduced in rmTBI mice vs. controls. Immunofluorescence for ADK (green) and GFAP (red) in the cerebral cortex (A,B) and the corpus callosum (C,D). Data were analyzed by one-way ANOVA followed by Tukey’s multiple comparison test. Statistical significance is indicated as follows: p < 0.05 (*), p < 0.01 (), p < 0.001 (***) n=4/group. Scale bar represents 50 µm.

**Supplementary Figure 4.** Activated microglia are increased at 72 h after rmTBI. Immunofluorescence for CD68 (green) and IBA1 (red) in the cerebral cortex (A,B) and the corpus callosum (C,D). Data were analyzed by one-way ANOVA followed by Tukey’s multiple comparison test. Statistical significance is indicated as follows: p < 0.05 (*), p < 0.01 (), p < 0.001 (***) n=4/group.

**Supplementary Figure 5.** Mice recover from sensorimotor deficits subsequent to rmTBI. A) The number of foot-slips recorded for mice on the horizontal ladder was tested and administered every 2 weeks after rmTBI. Data were analyzed by one-way ANOVA followed by Tukey’s multiple comparison test. Statistical significance is indicated as follows: p < 0.05 (*), p < 0.01 (), p < 0.001 (***), n=3 mice/group. B). Photograph of the horizontal ladder.

**Fig S6. Group differences in T1 (FLASH) and T2 (RARE) signal intensities were estimated using a two-sample t-test on a voxel-by-voxel basis across the whole brain.** A,D mTBI mice treated with vehicle vs sham mice, B,E. LIF-treated mTBI mice vs sham mice and C,F. vehicle-treated mTBI mice vs LIF-treated mTBI mice. Activated voxels have a threshold of P<0.05, corrected for multiple comparisons using a family-wise error control of 20-voxel contiguous cluster volume. Hot colors depict larger T1 or T2 intensity across the first group vs the second group. Sham mice (n=7), mTBI mice treated with vehicle (n=7) and mTBI mice treated with LIF (n=8). The arrow indicates the mTBI epicenter.

**Supplementary Figure 7.** Six remaining regions of interest which did not show significant FA change between mTBI treated with vehicle (blue) versus sham (green) or mTBI treated with both doses of LIF versus sham (brown and red).

**Supplementary Figure 8.** Six remaining regions of interest which did not show significant MD change between mTBI treated with vehicle (blue) versus sham (green) or mTBI treated with both doses of LIF versus sham (brown and red).

