## Supplementary figures and images for "Intranasal Leukemia Inhibitory Factor as a late-stage treatment for delayed white matter damage in concussive head injury"

### D'Mello Fig. S1

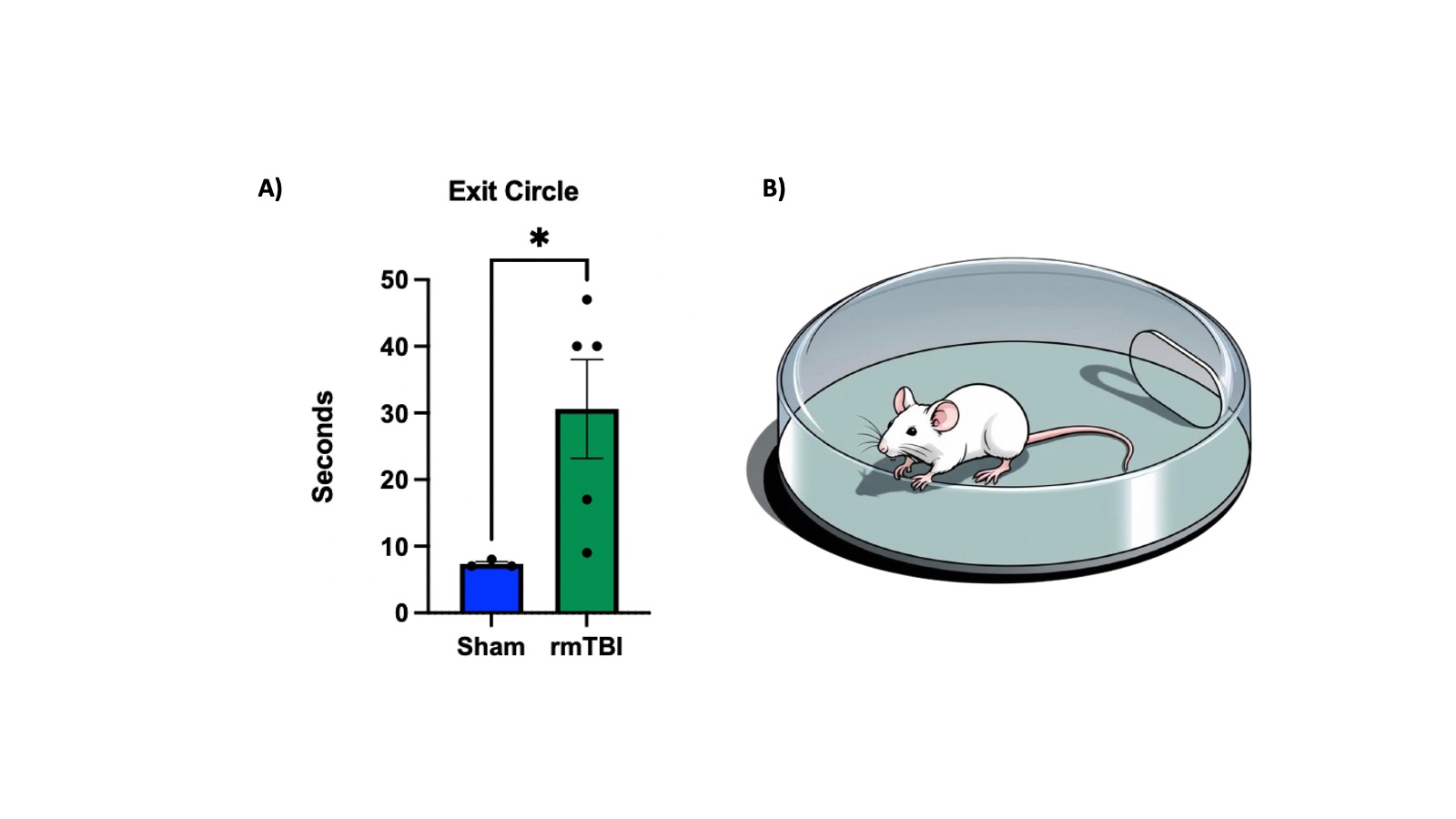

### D'Mello Fig. S2

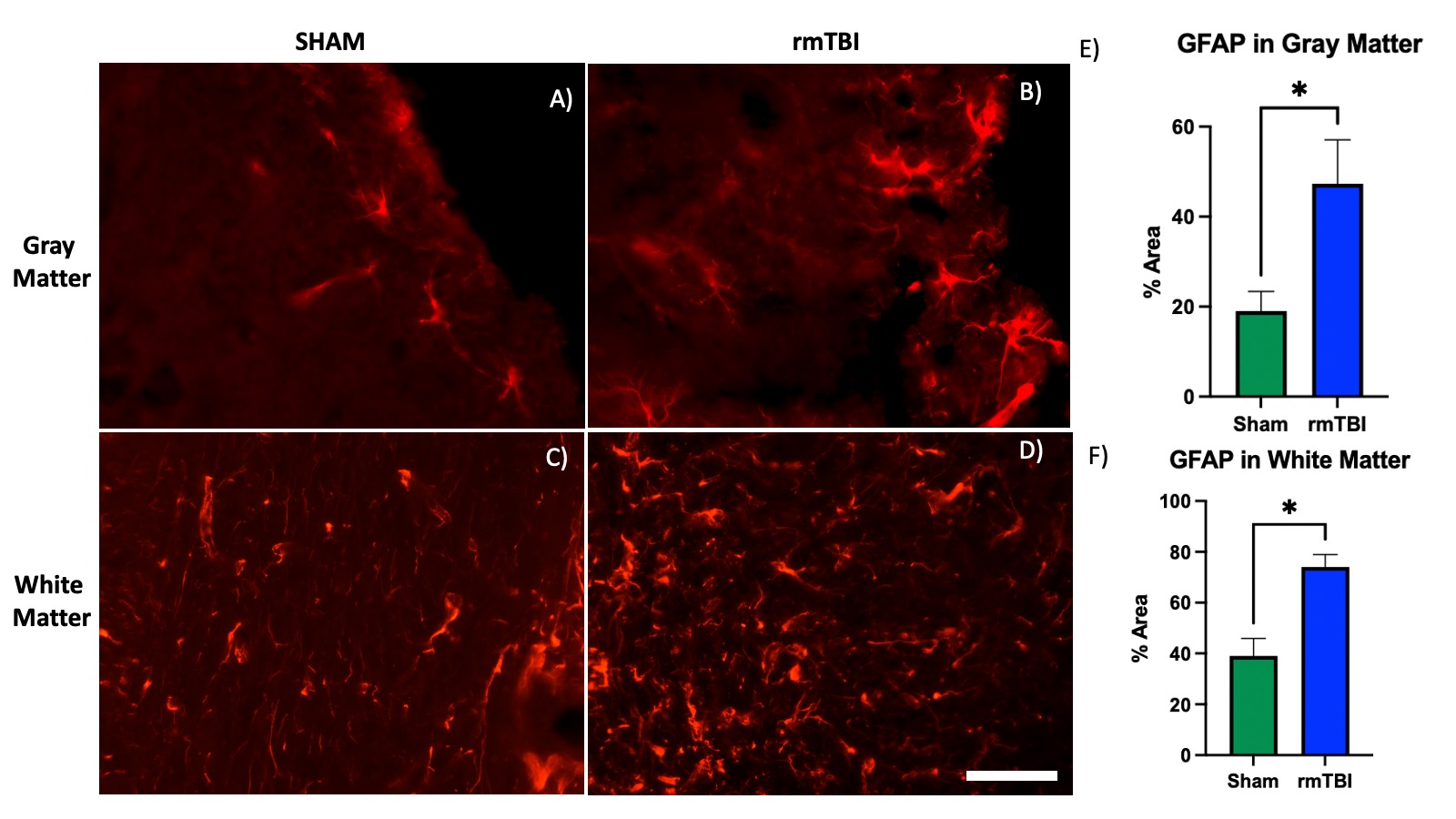

### D'Mello Fig. S3

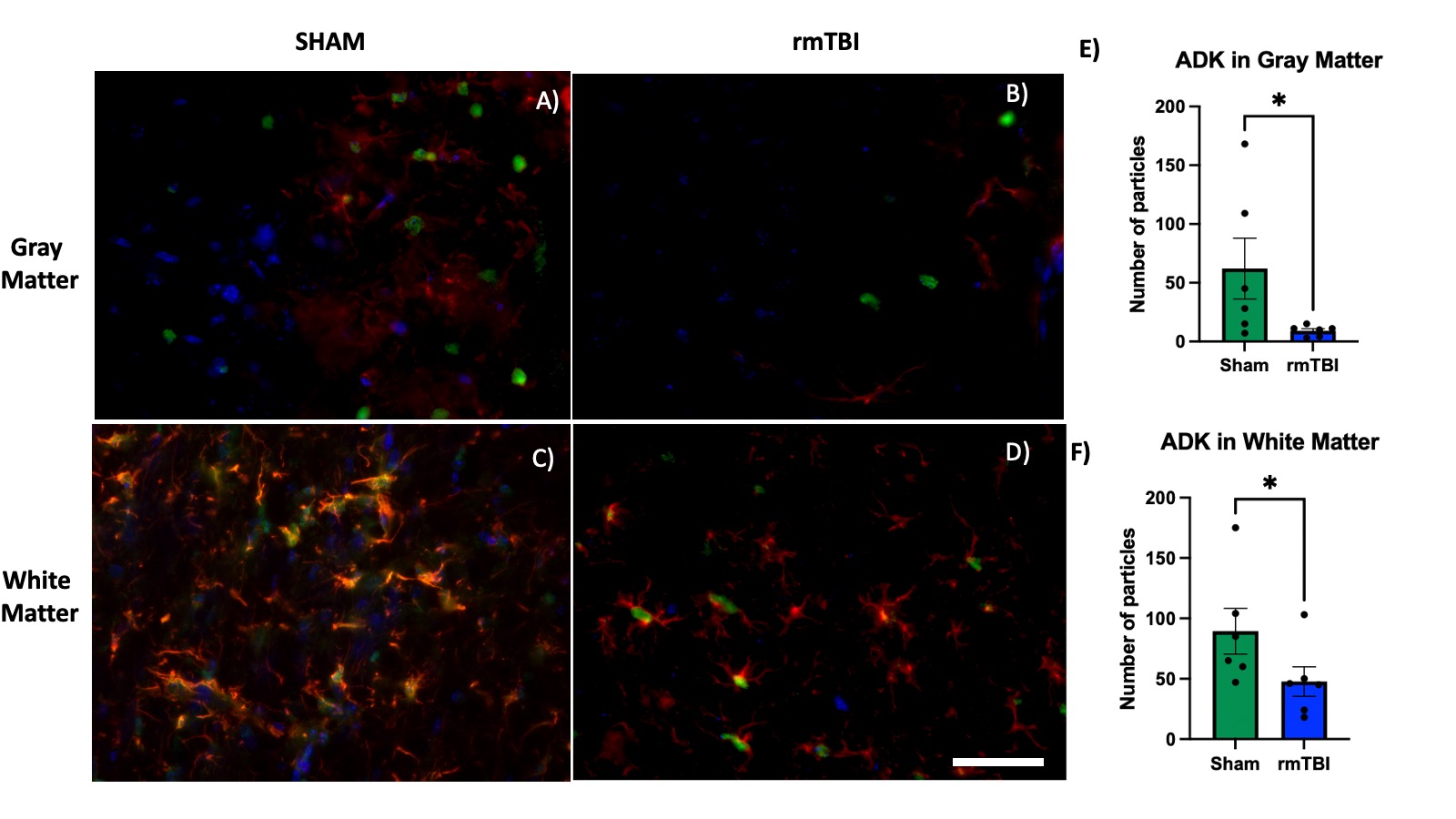

### D'Mello Fig. S4

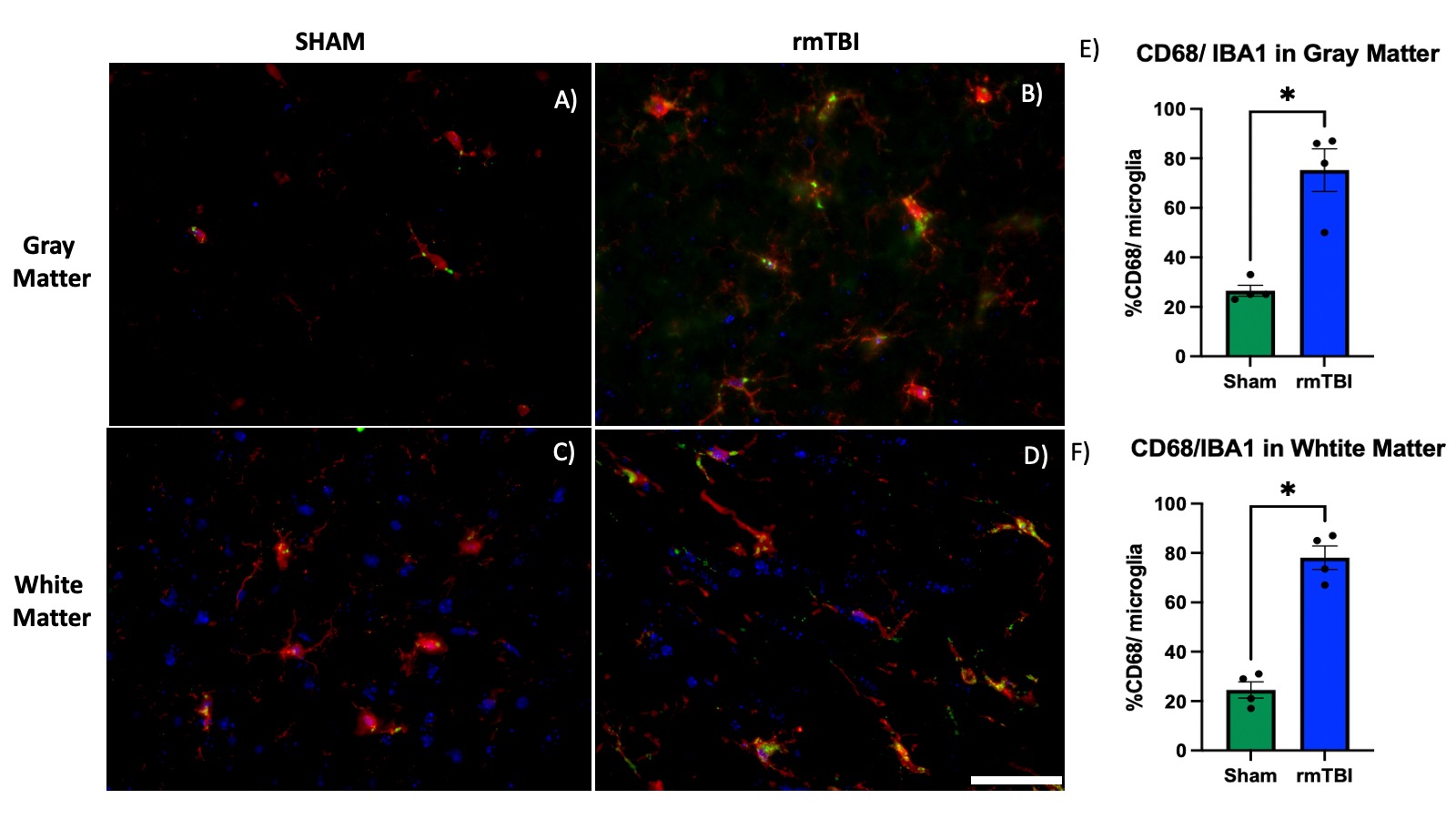

### D'Mello Fig. S5

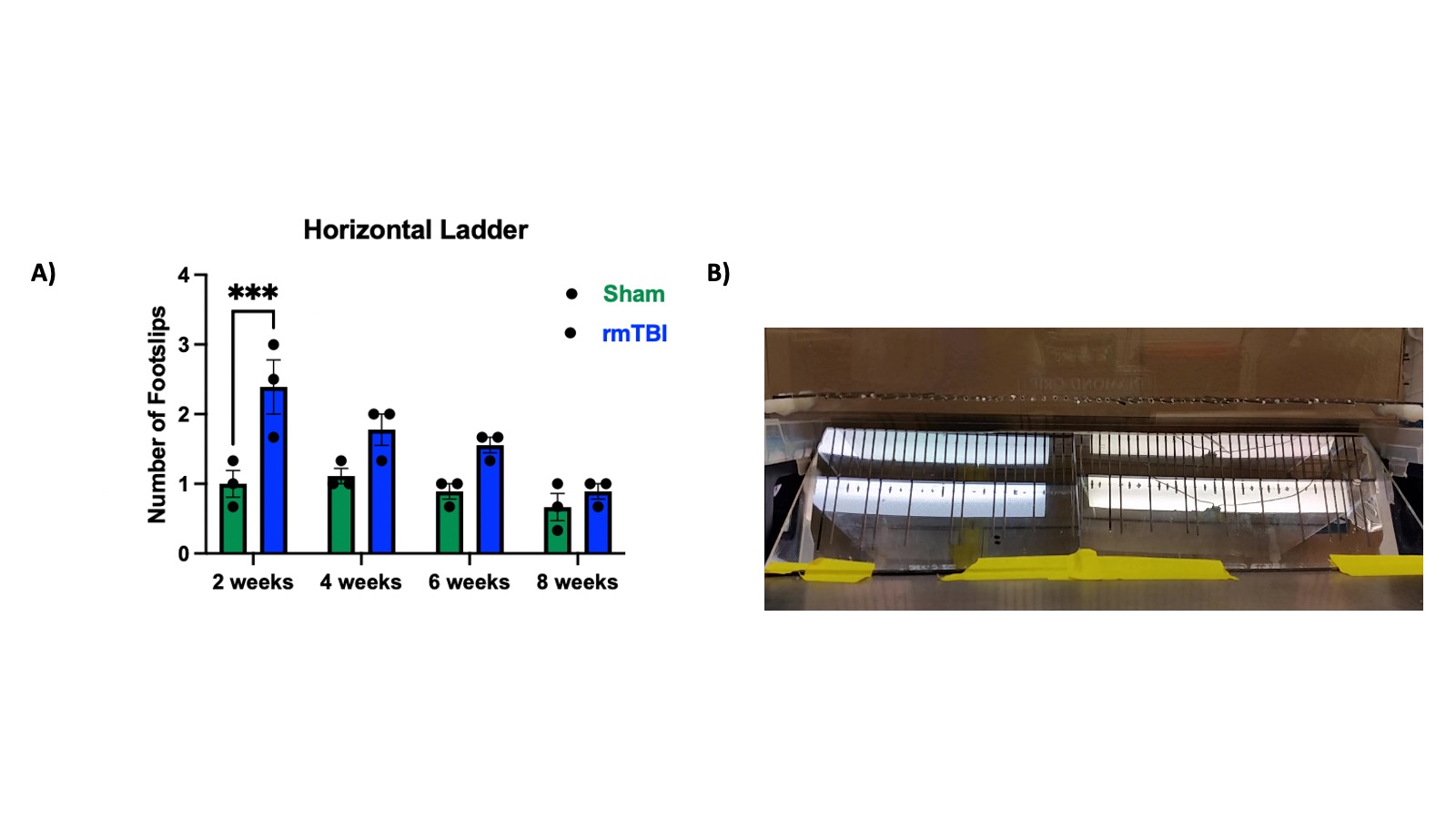

### D'Mello Fig. S6

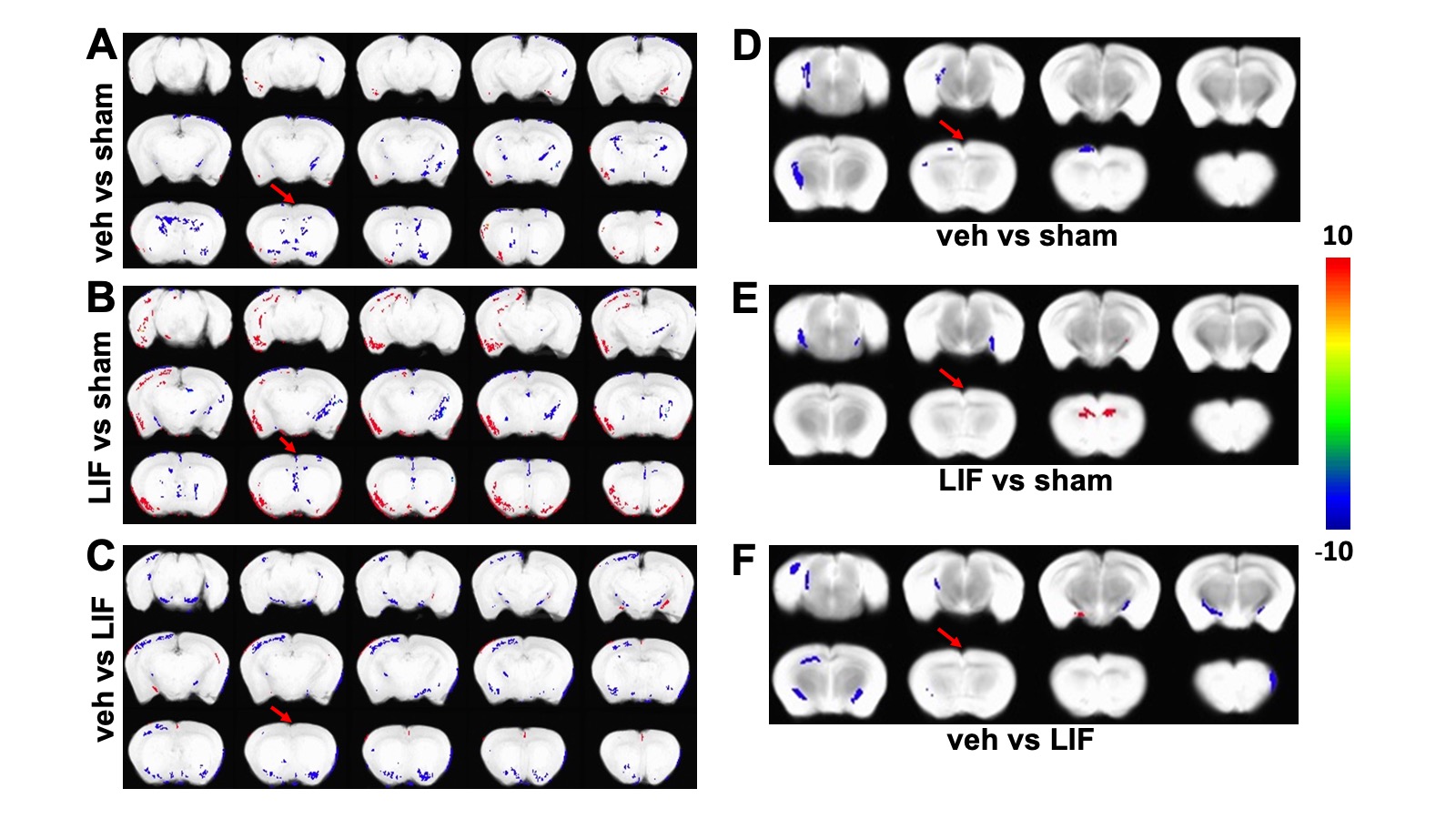

### D'Mello Fig. S7

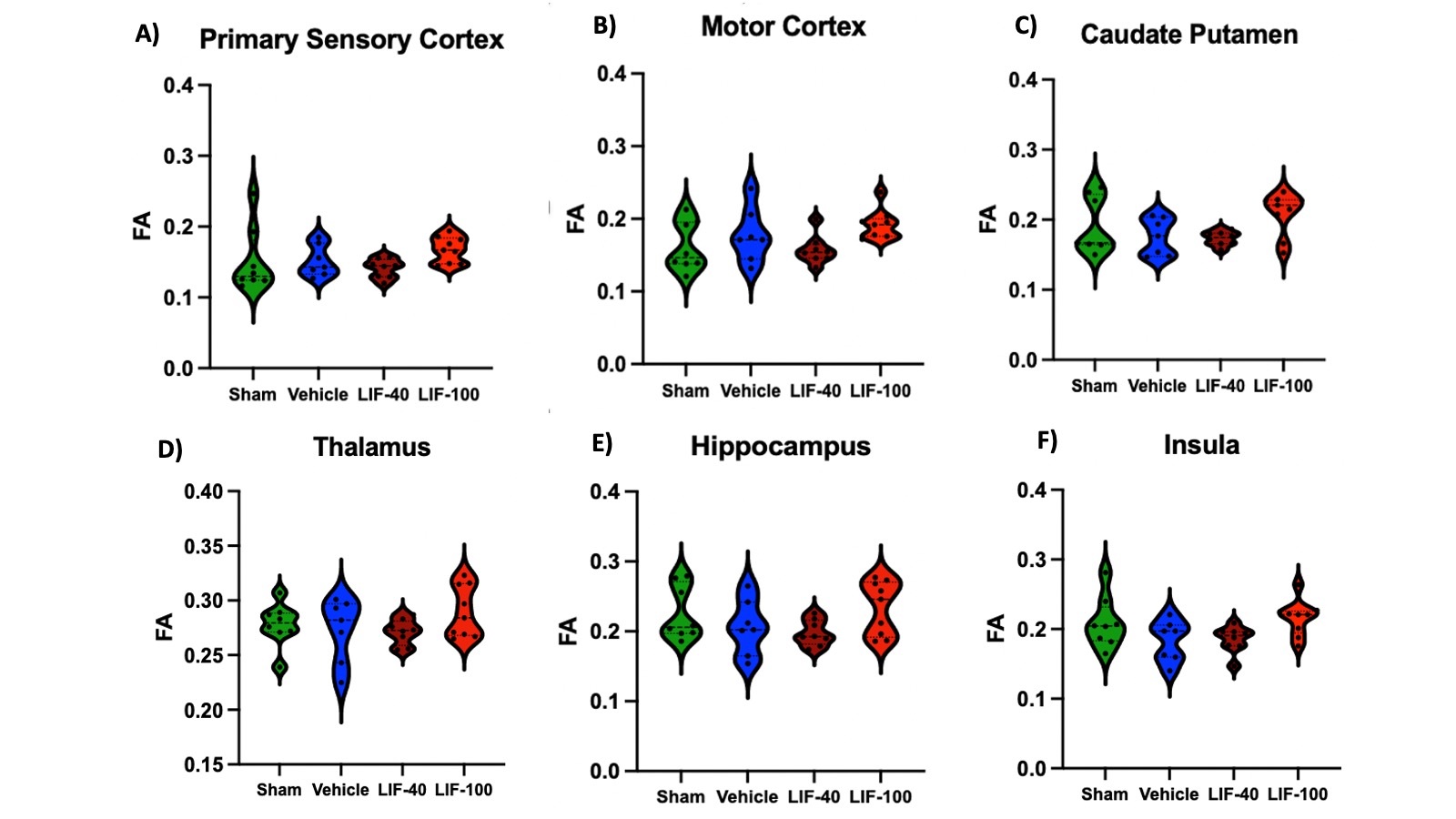

### D'Mello Fig. S8

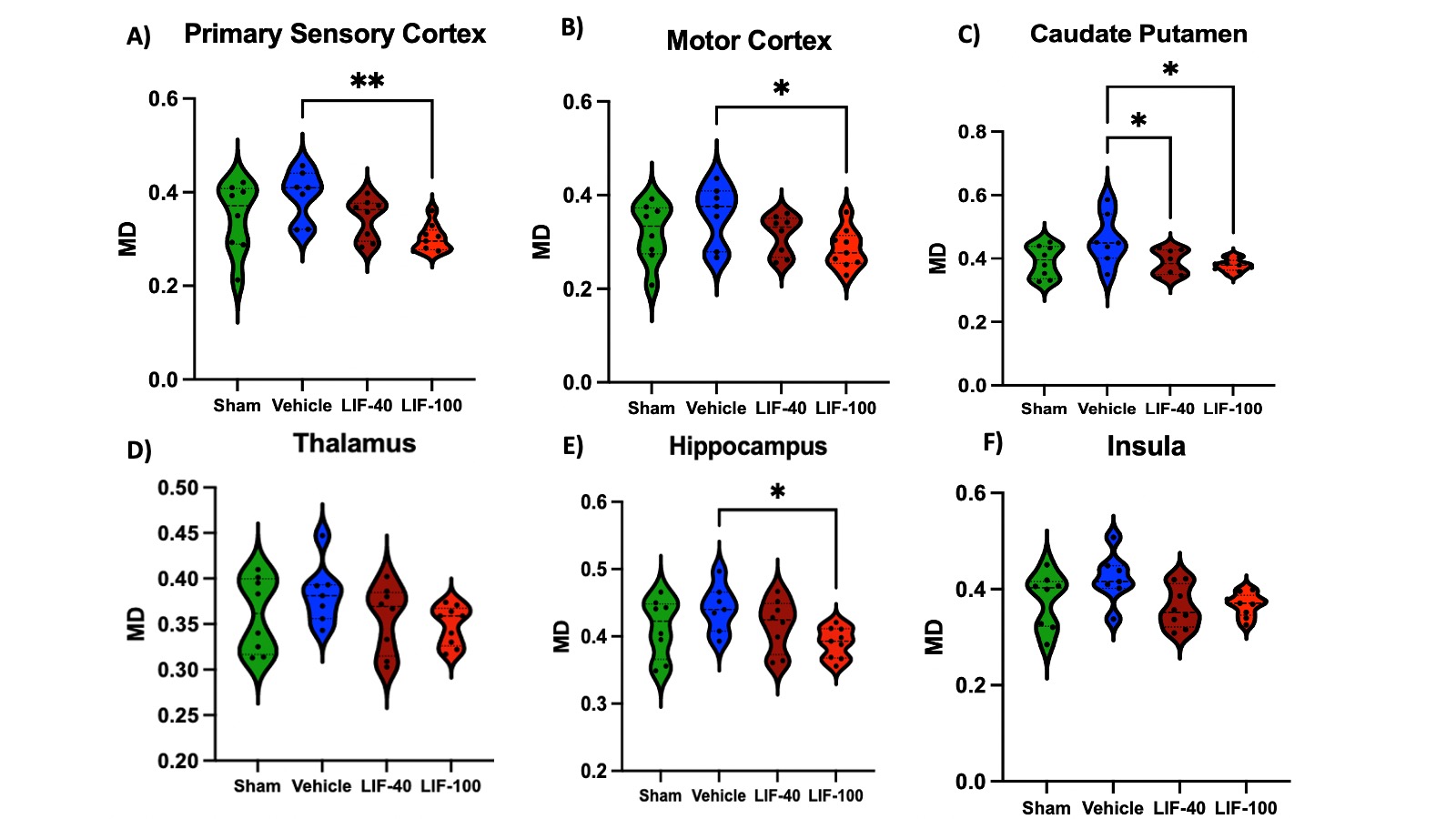
